# Multimodal, multifaceted Imaging-based Human Brain White Matter Atlas

**DOI:** 10.1101/2024.11.24.625092

**Authors:** Junchen Zhou, Wenxia Li, Shuo Xu, Huafu Chen, Jiao Li, Wei Liao

**Affiliations:** The Clinical Hospital of Chengdu Brain Science Institute, School of Life Science and Technology, University of Electronic Science and Technology of China, Chengdu 611731, P.R. China; MOE Key Lab for Neuroinformation, High-Field Magnetic Resonance Brain Imaging Key Laboratory of Sichuan Province, University of Electronic Science and Technology of China, Chengdu 611731, P.R. China

**Keywords:** multimodal white matter atlas, white matter function, multimodal 7-Tesla magnetic resonance imaging

## Abstract

Comparable to an atlas for navigating the Earth, a brain atlas outlines the brain’s neuroanatomical and functional landmarks. The tract-based brain’s white matter atlas (WMA) captures only tract boundaries, neglecting functionally relevant information, including emerging evidence on reliable neurodynamic detection in WM. Thus, a suitable WMA for functional investigations is needed. We delineated WM areas from coarse to fine-grained parcellations using local anatomical architecture and global functional dynamics using multimodal 7-Tesla magnetic resonance imaging in the Human Connectome Project. The multimodal WMA (MWMA) exhibited high between-subject consistency (69.1%) and test-retest reliability (93.1%), which aligned with the spatial pattern of chemoarchitectural homogeneity derived from open molecular imaging sources. Compared to a tract-based atlas, the MWMA better described the white matter’s cerebellum-association gradient axis, and precisely identified unique functional connectivity profiles of the corpus callosum. This improved MWMA will facilitate studies of WM functional organization, supporting cognition research and ultimately neuropathological vulnerability studies from a whole brain perspective.

## Introduction

The brain atlas, or parcellation-delineating spatial partitions, scaffolds the brain organization of structure and function(1, 2). The spatial arrangements of highly heterogeneous landscapes represent the specialized functions of different regions, and also allow investigations of their interactions and cooperations(3). Functional segregation and integration are complementary perspectives of human brain organization(4). Thus, identifying different specialized regions is fundamental to decoding the human brain. Early efforts to parcellate the mammalian brain using histological cytoarchitecture and myeloarchitecture(5), to recent *in vivo* structural and functional magnetic resonance imaging (MRI) data, have mostly involved cortical areas, and subcortical and cerebellar nuclei(6–12). Almost brain parcellations focus on the brain grey matter (GM)(13), which artificially ignores and excludes white matter (WM), hindering the development of the next generation human brain atlas(14).

WM is commonly referred to as the “other half brain,”(15, 16) and is composed of axonal projections from cortical GM neurons. Projections have historically been used to describe anatomical connections via WM tract pathways from diffusion MRI(17, 18). The tract-based WM atlas (WMA) provides the basis of structural brain connectomes(19, 20), but ignores functional information underlying the WM(21–23). WM has been found to have similar-sized venous veins to GM, and multiple functional activations involving the corpus callosum (CC), providing evidence for the potential neurophysiological origins of WM functional MRI (fMRI) signals(24, 25). Furthermore, relevant functional information of the WM was also suggested by our earlier researches, which ranged from investigations of neurogenetics(26), neurometabolism(27), neuroanatomy(28), neuro-electrophysiology(29), neurotopology(30), and neurobehavior(31) to neuropathology(32–34). Combining several features for parcellation studies can yield more information, because distinct traits identify various sets of regional borders. Because distinct features identify different sets of regional borders, combining multiple features for parcellation can provide additional information. Consequently, regional borders consistent across numerous independent attributes can be more confidently identified(8). To create a new WMA based on multimodal, multifaceted features, we thus identified brain WM subregions.

The basic premise of the brain atlas is to identify either topographically discrete areas or distributed networks, notwithstanding conceptual disagreements(35, 36). Therefore, the primary use of functional connectivity-based parcellation has been in the clustering of spatial elements (voxels) so that intrinsic connectivity fingerprints within a group of voxels are as comparable as possible, yet not within other groups(3, 37–41). The resulting clusters represent different brain regions or regions of interest (ROIs). To date, relatively few studies have used clustering to identify local properties(1). Regarding the homogeneity of functional signals, integrated local-global brain parcellation has outperformed local and global approaches(42, 43). Thus, a promising route for future brain parcellations is to combine global network fingerprints with local architecture or function(44–46). We have recently developed morphometrical and functional similarity networks, which are based on feature similarity and converge multimodal, multidimensional indicators of local attributes(47–50). As a result, we anticipate that applying this hybrid model synthesizing local and global neuromarkers should extend the conventional tract-based brain WMA(51–53).

To conceptualize WM function, we synthesized resting-state fMRI and diffusion-weighted imaging (DWI) data from a large cohort of 101 healthy subjects using a high field 7T scanner from Human Connectome Project. We used a two-level (individual and group levels) clustering technique to produce different atlas resolutions, when considering how inter-individual variability might impact the final group-level atlas. Furthermore, we assessed the test-retest reliability of the generated multimodal WMA (MWMA), and the resilience of MWMA to various disturbances. Then, using MRI features similarity transitions (macro level), and neurotransmitter similarity transitions (molecular level) derived from open molecular imaging sources, the validity of MWMA borders was evaluated. To test whether the MWMA better captured the functional topological organizations in WM, we compared the distance-controlled boundary coefficient (DCBC) and the dispersion of functional gradients between the MWMA and JHU-DTI-WMPM Type-I atlas(54). Finally, the functional connectivity patterns of the identified subregions in the CC further validated the applicability of MWMA in functional studies. The study design is shown in Fig. 1.

**Figure 1.**
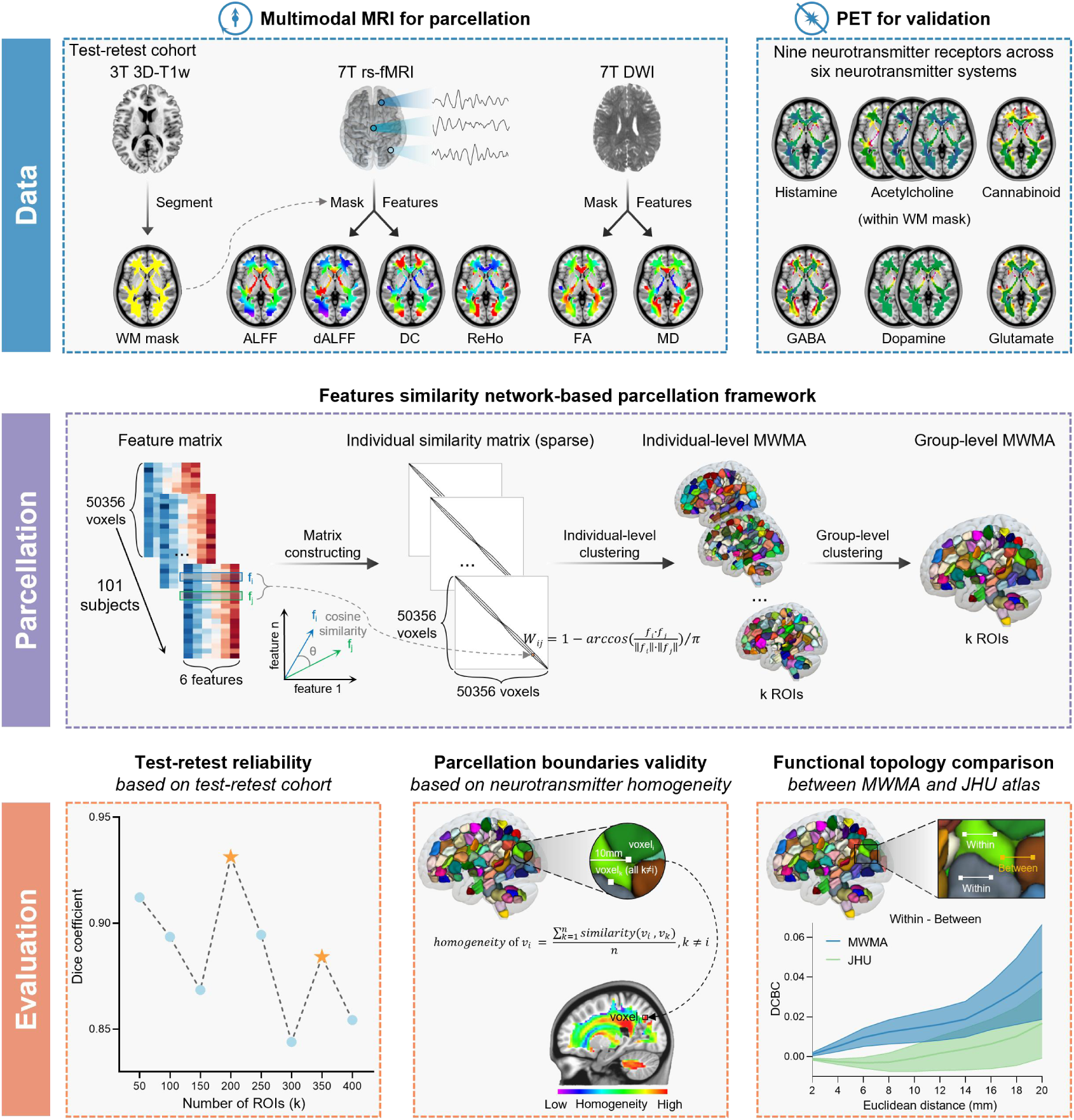
Study design. We parcellated the white matter (WM) based on the test-retest cohort (four rs-fMRI 16 min-scans, two scans per day) of 7-Tesla magnetic resonance imaging (MRI) data from the Human Connectome Project. The 3-dimensional T1-weighted images were used to generate the group-level WM mask. Six imaging features were extracted to construct a multifaceted similarity network, including the amplitude of low frequency fluctuations (ALFF), dynamic ALFF (dALFF), degree centrality (DC), and regional homogeneity (ReHo) based on the rs-fMRI images, and fractional anisotropy (FA) and mean diffusivity (MD) based on the diffusion-weighted imaging data within the group-level WM mask. To validate the WM atlas (WMA), we constructed neurotransmitter similarity transition based on nine different neurotransmitter receptors and transporters (D_2_, DAT, ɑ_4_β_2_, VAChT, M_1_, GABA_A/BZ_, mGluR_5_, H_3_, and CB_1_) across six different neurotransmitter systems (dopamine, acetylcholine, GABA, glutamate, histamine, and cannabinoid) collected by positron emission tomography analyses. Next, to integrate the six multifaceted MRI features into the global network, we constructed the individual network based on the cosine similarity of multifaceted features for each subject. Then, the individual network was submitted to the two-level spectral clustering procedure to generate the individual-level and group-level multimodal WMA (MWMA). To measure the MWMA’s test-retest reliability between the first and second scans, we calculated the Dice coefficient between parcellations. The local peak of the Dice coefficient indicated the potentially optimal resolution of the MWMA. In addition, given that voxels lying at the region of interest (ROI) boundary were expected to show lower homogeneity, we mapped MRI features and neurotransmitter similarity transitions to verify the ROI boundaries of the MWMA. Finally, to test whether the MWMA better captured the functional information of WM than the JHU-DTI atlas, we computed the distance-controlled boundary coefficient.

## Results

### MWMA based on multimodal, multifaceted features similarity networks

To obtain the MWMA based on multimodal MRI data, we first constructed multifaceted features similarity matrices at individual level (Supplementary Fig. 1). After performing the group-level spectral clustering, we clustered the whole brain WM into 50 to 400 ROIs with 50 intervals (Fig. 2A). As the MWMA resolution increased, the variability of ROI size across all ROIs decreased, and the features homogeneity of each ROI increased (Supplementary Fig. 2), indicating that the atlas resolution influenced the interpretation of the resulting MWMA. A coarse atlas might produce broad, over-generalized ROIs, whereas a fine-grained atlas produced a scattered MWMA map. For example, the 50-ROI MWMA divided the CC into four subregions (Supplementary Fig. 3), whereas previous parcellations of the CC suggested six to seven anatomically or functionally distinct subunits(55–57). In addition, the CC was the principal interhemispheric or commissural pathway in the brain, and the 400-ROI MWMA divided the CC into two hemispheres (Supplementary Fig. 3). Therefore, the 200-ROI MWMA may be a good compromise between the two extremes.

**Figure 2.**
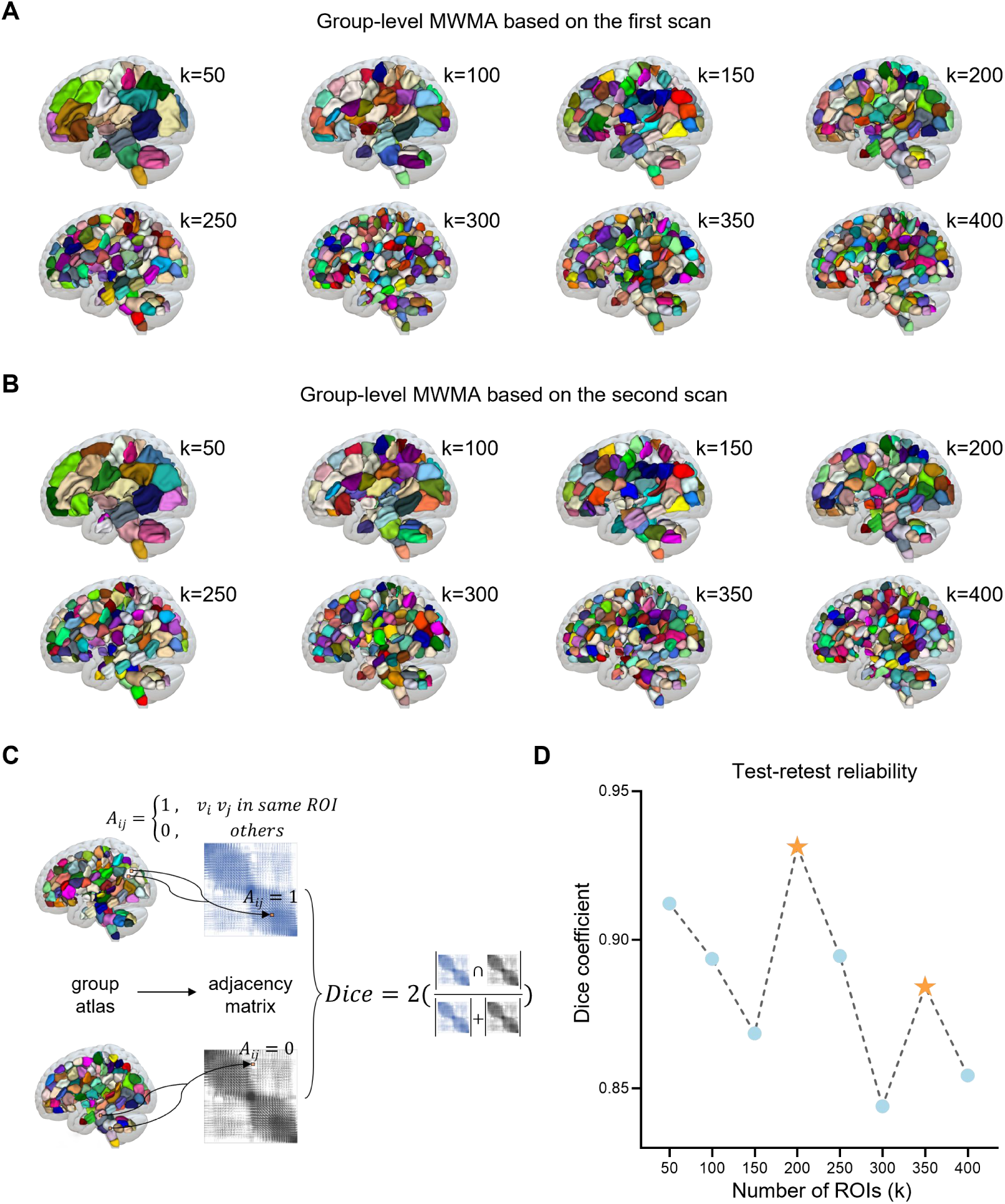
Multiple resolution multimodal white matter atlases (MWMAs) and their test-retest reliabilities. **(A)** The group-level MWMA with multiple resolutions based on the first scan [all 3-dimensional (3D) atlas images were visualized using the ITK-SNAP software(87)]. **(B)** The group-level MWMA with multiple resolutions based on the second scan. **(C)** Illustration of the MWMA Dice coefficient calculation was obtained from the first and second scans. The 3D group-level MWMA was converted into the adjacency matrix by setting the A_ij_ = 1, if the voxel_i_ and voxel_j_ were parcellated into the same region of interest (ROI). The Dice coefficient was then computed as twice the number of elements in the intersection of A_first scan_ and A_sceond scan_ divided by the sum of the number of elements in A_first scan_ and the number of elements in A_sceond scan_. **(D)** The test-retest reliability of the group-level MWMA. The resolution with the local Dice coefficient peak was chosen as the potentially optimal clusters. The stars represent the higher local Dice coefficient.

To measure the test-retest reliability of MWMAs between the first and second scans, we also parcellated the WM mask into a series of ROIs based on the second scan (Fig. 2B), which was the same as the first scan, and then calculated the Dice coefficient for each MWMA resolution (Fig. 2C). Although clustering similarities naturally decreased with larger numbers of ROIs, measuring the test-retest reliabilities facilitated the identification of highly stable solutions as local peaks on the graph (stars on the Fig. 2D). The parcellation maps were strikingly similar across the test-retest cohort.

Furthermore, the most stable multimodal parcellations were 200 ROIs (Dice coefficient = 0.93, Fig. 2D). The high-level test-retest reliability of the MWMA indicated that the 200-ROI MWMA might reflect the WM areal pattern of typical subjects in the healthy young adult population. All subsequent analyses were therefore based on the group-level 200-ROI MWMA.

In this MWMA resolution, some ROIs showed slightly different shapes and sizes across some individual areas, such as the frontal areas (Supplementary Fig. 4). However, the 200-ROI atlases were substantially similar across subjects [Dice’s coefficients (mean ± std) = 0.691 ± 0.069] (Supplementary Fig. 4), suggesting that this resolution of the MWMA may lead to minimal noisy estimates of a common multifaceted architecture.

We further evaluated the resilience of the 200-ROI MWMA to perturbations. We regenerated the 200-ROI MWMA using K-fold cross-validation. To generate a tiny and a relatively large perturbation, we performed 101-fold cross-validation (i.e., leave-one-subject-out) and 10-fold cross-validation, respectively. After repeating the clustering procedure 101 times, i.e., leaving one subject out of the current analysis, we found that the similarity of 200 parcels between the validation and primary atlas was at a high level, with the Dice coefficient ranging from 0.948 to 0.980 (mean ± std = 0.970 ± 0.007, Supplementary Fig. 5). Furthermore, using 10-fold cross-validation analyses, we repeated the procedure 50 times and found that validation parcellations exhibited a high Dice coefficient with the primary parcellation across 50 times (mean ± std = 0.960 ± 0.003, Supplementary Fig. 5), indicating the stability of the 200-ROI MWMA.

### Boundaries of the MWMA validated by multifaceted MRI features and neurotransmitters similarity transitions

The reproducible MWMA provided a global view of the multimodal, multifaceted atlas in WM. Next, we aimed to characterize the MWMA at a finer local scale. Given that voxels lying at the ROI boundary were expected to show lower homogeneity with their surrounding voxels (Fig. 3A), we mapped the multifaceted MRI features and neurotransmitter similarity transitions to identify the potential boundaries between ROIs as a way to support the validity of the MWMA(58) (Fig. 3B).

**Figure 3.**
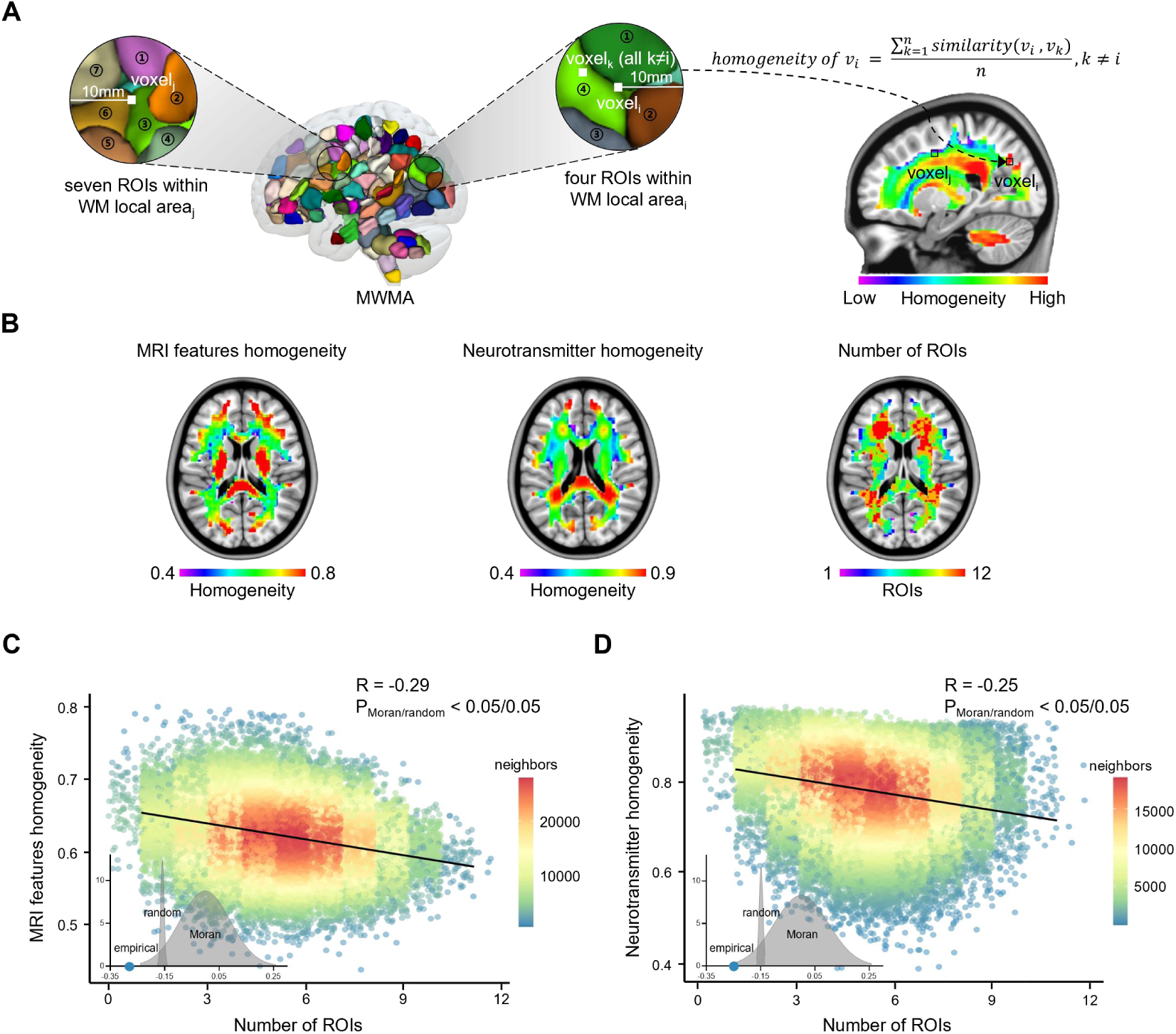
Validation of the multimodal white matter atlas (MWMA) by mapping the magnetic resonance imaging (MRI) features and neurotransmitter similarity transitions. **(A)** The illustration of MRI features/neurotransmitter homogeneity calculation and definition of the number of regions of interest (ROIs) in the voxel’s local WM area. The voxel’s local WM area was defined as the spherical center with a radius of five voxels (10 mm). For each voxel, we computed the cosine similarity of MRI features/neurotransmitters between this voxel and voxels within the voxel’s local area. The voxel’s homogeneity was the averaged similarities. **(B)** The spatial patterns of the MRI features homogeneity, the neurotransmitters homogeneity, and the number of ROIs for each WM voxel. **(C)** There was a significant negative correlation between the MRI features homogeneity and the number of ROIs across the whole brain WM. **(D)** There was a significant negative correlation between the neurotransmitter homogeneity and the number of ROIs across the whole brain WM.

For multimodal MRI data, we found a strong negative correlation (*R* =-0.29, *P*_moran_ < 0.05, Fig. 3C) between the number of ROIs of the voxel’s local areas, which was defined as the voxel as the spherical center with a radius of 10 mm and the MRI features showing homogeneity across all WM voxels, which supported the clustering-based parcellation to ROIs. To test whether this relationship was nonrandom, we constructed random atlases with 200 ROIs based on a random network (Supplementary Fig. 6), i.e., swapping each edge of the trivial similarity matrix twice while preserving the degree distribution and connectivity. We then evaluated the parcellation boundary validities of these random atlases. The correspondence between the number of ROIs and MRI features homogeneity in the voxel’s local area was more significant in the empirical relationship than in the null model (*P*_random_ < 0.05, Fig. 3C, inserted panel).

Next, we investigated the validity of the MWMA boundary based on the neurotransmitter receptors and transporter maps from open molecular imaging sources(59). The whole brain WM profile of neurotransmitter receptor densities was constructed by collating PET images from a total of nine different neurotransmitter receptors and transporters across six different neurotransmitter systems, including dopamine (D_2_ and DAT), acetylcholine (ɑ_4_β_2_, VAChT, and M_1_), GABA (GABA_A/BZ_), glutamate (mGluR_5_), histamine (H_3_), and cannabinoid (CB_1_). We masked all PET tracer maps using the group-level WM mask and z-score across the WM voxels, and calculated the cosine similarities of receptor/transporter fingerprints between pairs of voxels to measure the likelihood that two voxels were similarly regulated by endogenous or external input, which was referred to as “neurotransmitter similarity.” We also found a strong negative correlation between the number of ROIs in the local areas of voxels and neurotransmitter homogeneity across all WM voxels (*R* =-0.25, *P*_moran_ < 0.05, Fig. 3D). This relationship was not influenced by the random network-based parcellation procedure (*P*_random_ < 0.05, Fig. 3D, inserted panel), supporting the molecular substrate of the 200-ROI MWMA.

### Accurate functional and topographical organizations reflected by the MWMA

To estimate the validity of the generated MWMA, we computed the distance-controlled boundary coefficient (DCBC)(60, 61) for a pair of voxels. The fundamental tenet of the DCBC was that for any given distance in the brain, the functional connectivity (FC) strength between voxels within a ROI (within ROIs) should be greater than between voxels in separate ROIs (between ROIs) when a border correctly divided two functionally homogeneous ROIs (Fig. 4A). We calculated the DCBC (within-between) values between the pairs of WM voxel at spatial distances from 2 to 20 mm with a step of 2 mm. The DCBC values increased as the spatial distances between pair voxels increased; a high DCBC value indicated a high within-ROIs homogeneity and between-ROIs heterogeneity. Furthermore, we compared the MWMA with the existing tract-based atlas, i.e., the JHU-DTI-WMPM Type-I atlas. We found that the MWMA was better for presenting WM functional connectivity, when compared to the tract-based JHU-DTI atlas at each spatial distance between voxels (*P*_FDR_ < 0.05, Fig. 4B).

**Figure 4.**
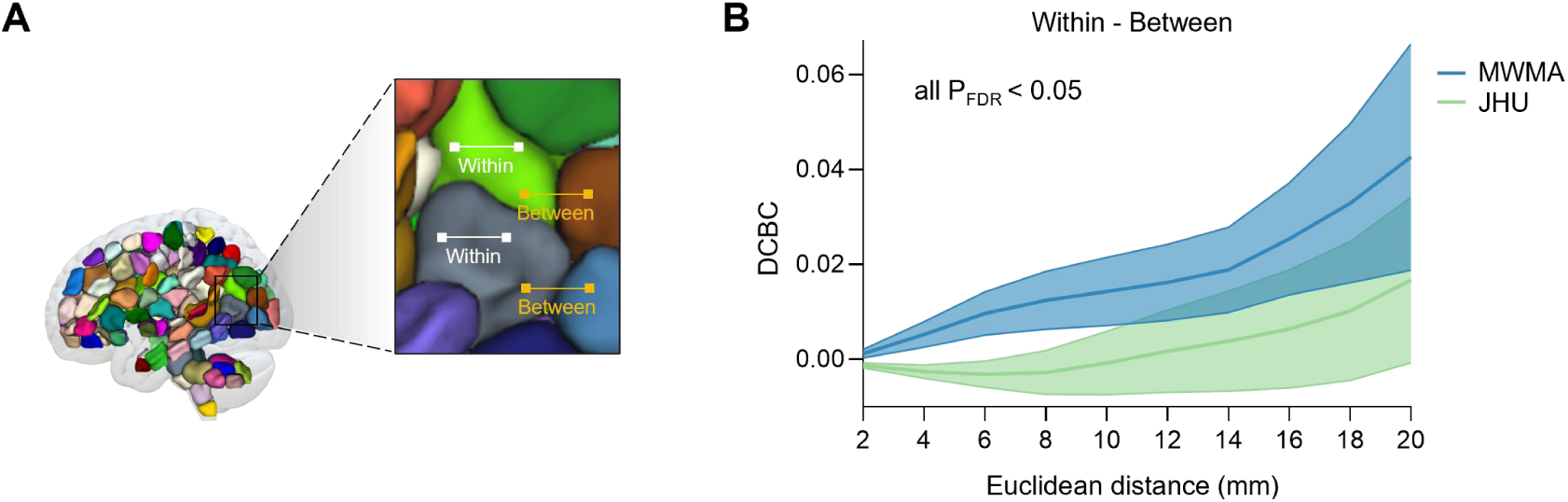
Validation of the multimodal white matter atlas (MWMA) using the distance-controlled boundary coefficient (DCBC). **(A)** Illustration of a DCBC calculation. The DCBC was calculated as the difference between the mean within regions of interest (ROIs) functional connectivity (FC) values and the mean between-ROIs FC values. **(B)** The comparison of the DCBC between the MWMA and the tract-based JHU-DTI atlas at a range of 2−20 mm with a step of 2 mm. The MWMA exhibited a better DCBC performance than the tract-based JHU-DTI atlas at each spatial distance (all *P_FDR_* < 0.05), indicating the more accurate capture of functional boundaries.

In addition to clear functional boundaries, the MWMA also preserved the topographical organization of the WM FC. To characterize the complex WM connectivity pattern, we mapped the FC onto the low dimensional representations as gradients, characterizing the topographical organization of the functional connectomes. We divided the 200 ROIs into seven functional networks based on a previous study(62), including a Default Mode Network (DMN), Frontoparietal Network (FPN), Sensorimotor Network (SMN), Visual Network (VN), Auditory Network (AN), Cerebellum Network (CBN), and Brainstem Network (BSN) (Fig. 5A). We determined the patterns of gradient spectrum for WM FC based on the MWMA and JHU-DTI atlases(27) (Fig. 5B and C, left panel). Furthermore, the dispersion of WM functional gradients provided a new means of studying the segregation and integration of functional brain networks in a multi-dimensional connectivity space(63, 64). We anticipated that if the MWMA captured topological organization better than the JHU-DTI atlas, the MWMA should exhibit lower dispersion of functional gradients than the JHU-DTI atlas. As shown in Fig. 5D, our MWMA resulted in less global dispersion (*P* < 0.05) and within-network dispersion in the FPN, SMN, and AN (all *P_FDR_* < 0.05). Therefore, our MWMA provided an alternative view to understanding the functional connectome of WM by reflecting the characteristics of the FC.

**Figure 5.**
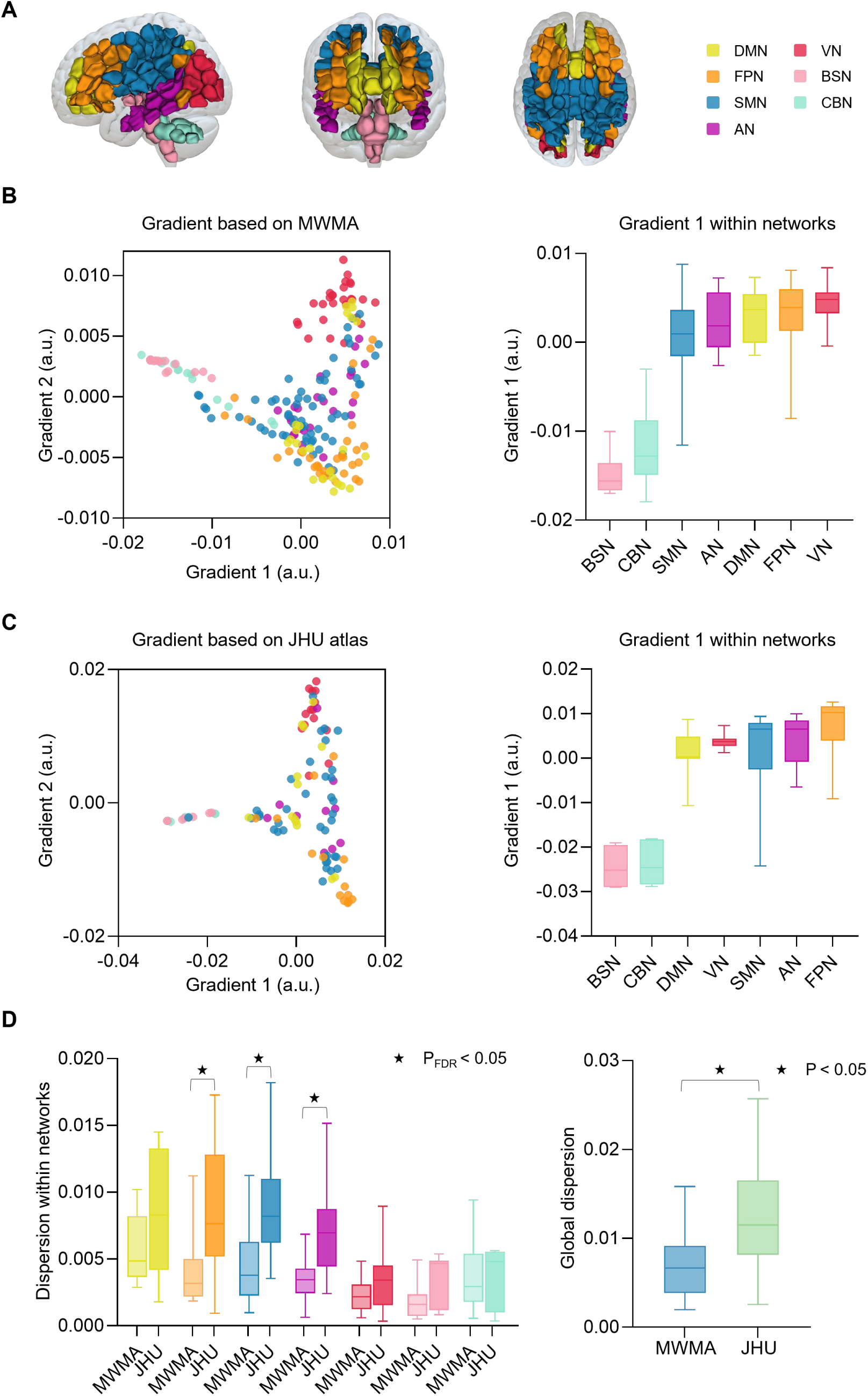
Validation of multimodal white matter atlas (MWMA) by mapping WM functional gradients. **(A)** The 200 regions of interest of the MWMA were divided into seven functional networks, including Default Mode Network (DMN), Frontoparietal Network (FPN), Sensorimotor Network (SMN), Visual Network (VN), Auditory Network (AN), Cerebellum Network (CBN), and Brainstem Network (BSN). **(B)** The functional gradient pattern of the WM was based on the MWMA. The first two functional gradients were scattered in a typical triangular distribution. The first gradient was anchored at the two extreme ends, the BSN and VN. **(C)** The functional gradient pattern of white matter was based on the JHU-DTI atlas. The first two functional gradients were also scattered in a typical triangular distribution, while the first gradient was anchored at the two extreme ends, the BSN and FPN. **(D)** Comparisons of dispersions of functional gradients between the two atlases. The MWMA showed a lower global dispersion, and a lower within-network dispersion in the FPN, SMN, and AN than the JHU-DTI atlas, indicating the more accurate representation of WM functional topographical organizations.

### Functional connectivity patterns of the CC derived from the MWMA

After elucidating the reproducibility and validity of the MWMA, we next determined the FC patterns of the identified subregions in detail to validate the applicability of the MWMA in functional studies. The CC, the largest WM fiber bundle connecting the brain hemispheres, has been recognized in several subregions using the FC technique(65). Here, we parcellated the CC into six subregions in the MWMA (Fig. 6). In our atlas, CC 1 and CC 2 were mainly located on the rostrum of CC; CC 3 mostly belonged to the midbody; and CC 4, CC 5, and CC 6 were primarily located on isthmus and splenium. We then calculated the FC of each CC subregion, and found similar connectivity patterns between whole WM ROI-levels and voxel-levels (*P*_Moran_ < 0.05, FDR corrected), indicating the high functional homogeneity of parcellated regions. In addition, the FC across six CC subregions exhibited unique patterns. The subregions on the rostrum of the CC showed strong connections with DMN and FPN (Fig. 6A and B). The CC 3 strongly connected with cerebellum and brainstem networks (Fig. 6C), and the CC posterior regions functionally connected with FPN and VN, and also showed strong connections with DMN (Fig. 6D-F).

**Figure 6.**
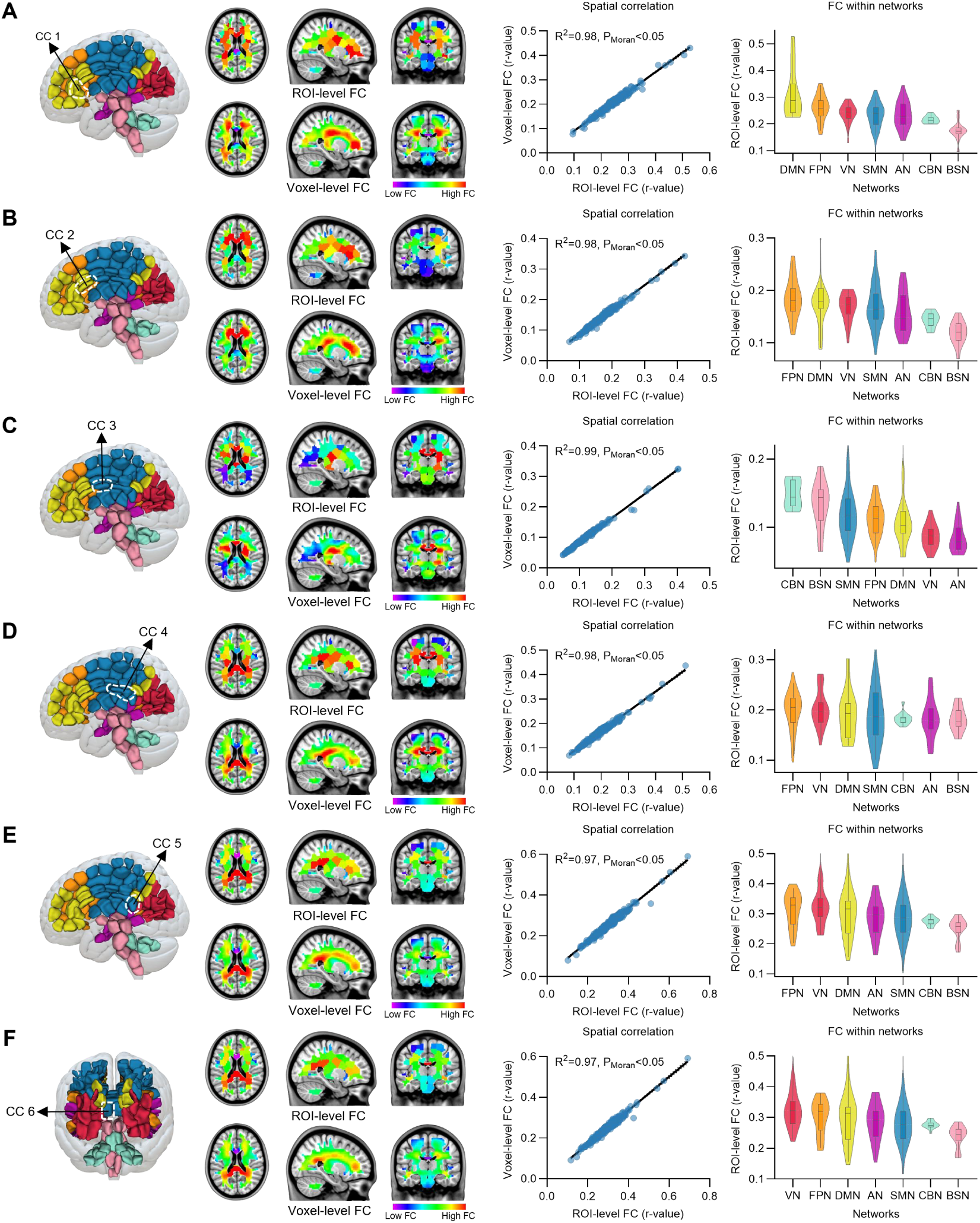
Functional connectivity (FC) pattern of subregions of the corpus callosum (CC). The CC was parcellated into the six subregions in the 200-region of interest (ROI) multimodal white matter atlas (MWMA). Each subregion showed a unique functional connectivity (FC) pattern with high reproducibility of the voxel-level FC pattern, indicating the applicability of the MWMA in functional connectivity studies. The first panel shows the location of each subregion of the CC. The second panel shows the whole WM FC patterns at the voxel-level and ROI-level. The third panel shows the spatial correlation between each CC subregion’s ROI-level and voxel-level FC patterns. The fourth panel shows the whole WM functional connectivity pattern of each subregion of the CC from the functional network perspective. The FC pattern was sorted by the median of the FC values within each network.

## Discussion

We have generated a robust human population-based MWMA using high resolution multimodal data from hundreds of test-retest cohorts based on Human Connectome Project subjects. The MWMA provided a new framework for human brain functional research, particularly whole brain connectome analyses, that overcame several drawbacks of previous parcellation schemes as follows: i) it established a neurobiologically valid brain parcellation scheme of the entire WM showing a coherent pattern of chemoarchitectural homogeneity; and ii) it provided more detailed characterizations of the WM FC profiles than the tract-based JHU-DTI atlas.

Due to increasing studies of WM functional activity measured by conventional fMRI, a new human brain WMA with a framework for integrating multimodal, multifaceted features is urgently needed. Consequently, many studies have used different MRI modalities to identify brain WM subregions or to identify functional systems of WM(27, 54, 58, 62). While there is no agreement on which modality or aspect of brain organization most represents brain WM organization, brain WM atlases are essential to further our understating of the human brain. This is because coordinate systems or anatomical landmarks are invalid markers of regional specialization(9). This need is met by the MWMA with a framework for integrating multimodal, multifaceted features, providing a whole brain WM parcellation into distinct subregions according to the multifaceted features of the similarity connectome. By leveraging the local similarity connectivity architecture, we identified the ROIs that were different from one another, while maintaining a high degree of internal homogeneity within ROIs(37). Furthermore, we used 7 Tesla (7T) MRI to probe fine-grained parcellation in this study, which can produce high resolution images with high signal-to-noise and contrast-to-noise ratios(66). Based on 7T functional MRI data, one study has used independicent component analysis to identify 30 resting-state networks within white-matter mask(44). Compared to this work, we provided a more fine-grained parcels based on multimodal MRI data. Combining multiple features might provide complemental information, because different features can distinguish different sets of areal boundaries(7, 8). Take together, our MWMA provided a robust foundation for investigating the intricate functional connections of WM.

By generating a reliable neuroanatomical map of human WM regions, the present study represented a significant advance relative to previous human brain WM atlases. Our approach showed that we could generate a highly reproducible MWMA using test-retest multimodal brain neuroimaging data. This MWMA was consistent with chemoarchitectural homogeneity. In addition, compared to the tract-based JHU-DTI atlas, our MWMA showed more accurate functional and topographical organizations. This improvement makes it possible to use spatial clusters that move beyond the traditional use of tracts combined with the JHU-DTI atlas label, to investigate WM functional information using fMRI studies. From a neuroanatomical perspective, the MWMA based on multimodal MRI neuroimaging provided certainty regarding whether any two functional neuroimaging studies resulted from the same WM regions.

In addition to confirming other differentiations from prior cytoarchitectonic and myeloarchitectonic maps, the MWMA also identified anatomical subregions that were previously not documented(67–69). In the MWMA, the CC has been divided into six subregions, but the precise parcellation of the human CC is controversial(55–57). However, the current parcellation identified specific functional connectivity for CC subregions, consistent with a previous functional map(65), which showed the anterior to posterior aspect of the CC. In addition, the posterior of the CC was divided into left and right symmetric parts, which was consistent with the third functional gradient map of the CC, possibly contributing to functional lateralization of the cerebral hemisphere(70). These findings increased our understanding of the role of the CC in interhemispheric information transmission, and highlighted the need for more research.

Although we provided a comprehensive multimodal data resource for mapping the human brain WM functional connectome, our current study still had limitations that need to be addressed by future research. Young adults were used to generate the MWMA, but it did not cover the whole human lifespan. Because brain development is critical for accurately mapping a fully functional connectome, including more cohorts of different ages will be necessary to fully construct a better functional architectures of WM.

Together, the current work provided an open-source MWMA for use in functional research on human brain WM. As the number of studies of WM functional activity increases, we hope that a multimodal, multifaceted WMA will provide investigators with a standardized atlas to better understand what WM functional information can reveal about the human brain.

## Materials and Methods

### Participants and MRI acquisition

Among the participants using the HCP S1200 release 7T MRI, a total of 174 participants out of the original 184 participants received complete diffusion-weighted imaging (DWI) and four rs-fMRI 16 min-scans (two scans per day), namely REST1_PA, REST1_AP, REST2_PA, and REST2_AP. The PA and AP represented posterior-to-anterior and anterior-to-posterior phase coding directions that were used in the gradient-echo EPI sequence. The rs-fMRI and DWI images were acquired on the 7T scanner (Siemens Healthcare, Germany) using a gradient-echo EPI sequence at 1.6 mm isotropic resolution and a spin-echo EPI sequence at 1.05 mm isotropic resolution, respectively. Specifically, the REST1_PA and REST1_AP scans were used for the main parcellation, whereas REST2_PA and REST2_AP scans were used to evaluate the test-retest reliabilities. In addition, the high resolution (0.7 mm isotropic voxel) structural images of T1-weighted (T1w) sequences were acquired on a customized Siemens 3T Connectome Skyra scanner (Siemens Healthcare, Germany) using a magnetization prepared rapid acquisition gradient-echo sequence (MPRAGE). More details about the imaging parameters have been described previously(71).

### MRI data preprocessing

For structural MRI images, the preprocessing pipelines were as follows: the first step was conducted based on a PreFreeSurfer pipeline; the second step was probabilistic segmentation of the T1w volume into WM, GM, and cerebrospinal fluid (CSF) using the FMRIB Software Library (FSL)(72). The final step downsampled the structural images to 2 × 2 × 2 mm^3^ when considering the computing efficiency and storage space based on our previous study(73).

Diffusion-weighted images were first preprocessed using the HCP Diffusion processing pipeline. Further preprocessing involved the following steps: first, only the low b-value DWI volumes (b = 1000) from the 7T multi-shell DWI dataset were included in this study(74). Second, the DWI images were downsampled to 2 × 2 × 2 mm^3^ for the dot product with the group-level WM mask. Third, the individual b0 images were registered with corresponding individual T1w images to yield the transformation matrix 1, and the individual T1w images were registered in the Montreal Neurological Institute (MNI) space to yield transformation matrix 2. Subsequently, these two transformation matrices were used to register the individual fractional anisotropy (FA) and mean diffusivity (MD) images in the MNI space.

Functional images were first preprocessed through the HCP fMRI Volume pipeline. Further preprocessing involved the following steps: first, 73 subjects were excluded due to a Root Mean Squared head position change over half a voxel’s width (0.8 mm) from the four rs-fMRI scans(75), leaving a cohort of 101 healthy young adults (59 females, age = 29.35 ± 3.08).

To minimize mixing signal components from the WM and GM due to partial volume effects, we performed subsequent processing of functional images within the WM mask. First, an individualized WM mask from each participant was obtained using a 50% threshold on the probability maps of WM (i.e., produced by probabilistic structural segmentation). Only voxels identified as WM across 80% of participants were included as part of the group-level WM mask. Second, the Harvard Oxford Atlas was used to remove subcortical nuclei from the group-level WM mask, including thalamus, caudate, putamen, pallidum, and accumbens(30, 31). Additionally, discrete marginal small areas in the group-level WM mask were excluded by identifying the connected component with less than 20 voxels in each 3D section. Consequently, the group-level WM mask comprised 50,356 voxels at 2 mm isotropic resolution (Supplementary Fig. 7).

After performing dot product between functional images and the group-level WM mask for each participant, we down-sampled the functional images to 2 × 2 × 2 mm^3^ to match the resolution of the structural images. Multiple regression of 12 rigid body motion parameters (six head motion parameters and six head motion parameters, one time point before), and the mean CSF signals were regressed-out from the functional images(75–77). Then, a band-pass filter of 0.01−0.1 Hz was performed to reduce the impact of non-neuronal factors(78). Finally, four rs-fMRI time series were demeaned, and then the REST1_PA and REST1_AP time series were concatenated, while the REST2_PA and REST2_AP time series were concatenated. Thus, the length of the rs-fMRI time series for each scan was 1,800 TRs.

### Construction of the individual multifaceted features similarity matrix

To construct the individual-level multifaceted features similarity network, we computed six functional and structural MRI features for each voxel in the WM, including the amplitude of low frequency fluctuations (ALFF), dynamic ALFF (dALFF), degree centrality (DC), regional homogeneity (ReHo), FA and MD.

Each voxel’s functional and structural metrics were divided by the WM mean value to normalize the global effects. For each subject, six functional and structural metrics were vectorized as a multifaceted feature matrix (50,356 × 6). Each feature vector (50,356 × 1) was z-normalized across voxels to account for variations in value distributions between the metrics.

The individual multifaceted feature similarity matrix W was constructed by calculating the cosine similarity of two given voxels’ feature vectors (1 × 6)(79). The cosine similarity was a non-negative similarity measure necessary for the spectral clustering algorithm(80). For each subject, the cosine similarity was computed between a given voxel and its 26 nearest neighboring voxels (spatial constraint). This resulted in a similarity matrix W with a sparsity of approximately 4/10,000(37). In addition, this spatial constraint produced contiguous ROIs and reduced the computational cost. The computation of the matrix W was defined by:

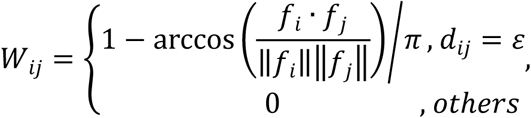

where, f_i_ and f_j_ denote the normalized feature vectors of two voxels i and j; the d_ij_ denotes the Euclidean distance of two voxels i and j; ε is equal to 1 or √2 or √3, ensuring that voxels i and j are only in each other’s 26 nearest neighborhoods. A symmetric, sparse, multifaceted feature similarity matrix W (50,356 × 50,356) was calculated for each subject.

### Two-level spatially constrained spectral clustering

The two-level spatially constrained spectral clustering (SCSC) had two main steps: the individual-level clustering based on the individual multifaceted similarity matrix W; and the group-level clustering based on the group averaged adjacency matrix A_group_.

### Individual-level spectral clustering

Individual-level spectral clustering involved the following steps: first, the individual multifaceted feature similarity matrix W was converted into the corresponding normalized Laplacian matrix L by:

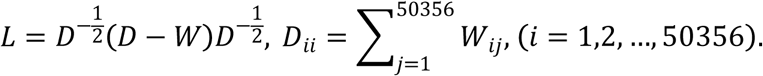

Second, spectral decomposition was performed on the L:

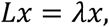

where λ is L’s eigenvalue, x is L’s eigenvector corresponding to eigenvalue λ. In practice, k eigenvectors corresponding to the k smallest eigenvalues were computed(80). Third, the column-pivoted QR factorization (CPQR) algorithm was performed to cluster the rows of k eigenvectors into k clusters, i.e., cluster the 50,356 voxels into k ROIs. The CPQR algorithm had higher computational efficiency without iterating and tuning parameters. In addition, this algorithm avoided random initialization, leading to poor clustering results compared to conventional clustering algorithms, such as k-means and multiclass spectral clustering algorithms(81). In this manner, 101 individual-level atlases with k ROIs were obtained (Supplementary Fig. 1). This study chose the number of ROIs k from 50 to 400 with a step of 50.

### Group-level spectral clustering

The steps of group-level spectral clustering were similar to those of individual-level spectral clustering. The only difference was that group-level spectral clustering was based on the group averaged adjacency matrix A_group_ instead of the individual multifaceted similarity matrix W. Specifically, group-level spectral clustering involved the following steps: first, the individual adjacency matrix A was obtained from the individual atlas according to the following rule: if two given voxels v_i_ and v_j_ were parcellated into the same ROI, the weight A_ij_ was set to 1:

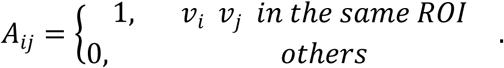

Second, individual adjacency matrices were averaged to obtain the group-averaged adjacency matrix A_group_ (Supplementary Fig. 1). Third, the group averaged adjacency matrix A_group_ was converted into the corresponding normalized Laplacian matrix L_group_. Then, spectral decomposition was performed on L_group_ to obtain k eigenvectors corresponding to the k smallest eigenvalues. The CPQR algorithm was then performed to cluster k eigenvectors. Finally, group atlases of 50−400 ROIs were obtained (Fig. 2A).

All ROIs were then assigned to WM functional networks. According to a previous study(62), 10 WM independent components (ICs) were identified based on the track-weighted dynamic FC, including Medial Sensorimotor (IC1), Medial Visual (IC2), Default Mode (IC3), Auditory (IC4), Lateral Sensorimotor (IC5), Frontoparietal (IC6, IC7), Premotor (IC8), Lateral Visual (IC9), and Vertical Occipital (IC10). The 10 ICs were merged into five functional networks: Default Mode Network (DMN, i.e., IC3), Frontoparietal Network (FPN, i.e., IC6 and IC7), Sensorimotor Network (SMN, i.e., IC1, IC5 and IC8), Visual Network (VN, i.e., IC2, IC9, IC10), and Auditory Network (AN, i.e., IC4). In addition, the cerebellum and brainstem were included in our analysis, and named as the Cerebellum Network (CBN) and Brainstem Network (BSN). Subsequently, seven WM functional networks were included in our analyses. According to the position of the centroid of each ROI belonging to a functional network, all ROIs were assigned to seven functional networks (Fig. 5A).

### Evaluation of the WMA

#### Test-retest reliability of the group-level atlas

To measure the test-retest reliability of the WMA, we calculated the Dice coefficient between atlases based on the first and the second scans at all resolutions from 50 to 400 ROIs. Clustering similarities decreased as the number of ROIs increased, so the potential optimal number of ROIs was determined by the local peak of the Dice coefficient(36, 58). The Dice coefficient was calculated by:

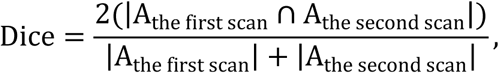

Where the A_the first scan_ and A_the second scan_ are adjacency matrices of atlases based on the first scan and the second scan, respectively. The Dice coefficient was categorized generally as follows: low (0−0.19), low moderate (0.20−0.39), moderate (0.40−0.59), moderately high (0.60−0.79), or high (0.8−1.0)(82).

#### Between-subjects atlas consistency

The consistencies of between-subjects atlases were assessed by measuring the Dice coefficient between the individual 200-ROI atlas and group-level 200-ROI MWMA based on the first scan.

#### Resilience of the atlas to perturbations

The robustness of the group atlas was evaluated by k-fold cross-validation (CV). In this study, 101-fold CV and 10-fold CV were performed. In each turn of the 101-fold CV, one subject was excluded. The remaining 100 subjects were submitted to the two-level SCSC procedure, resulting in a 100-subject group-level atlas. Then, the Dice coefficient between the 100-subject group-level atlas and the MWMA (101-subjects group-level atlas) was calculated.

In each turn of the 10-fold CV, 101 subjects were randomly divided into 10-fold, where 9-fold contained 10 participants, and 1-fold contained 11 participants. One-fold was randomly excluded, and the remaining subjects were submitted to the two-level SCSC procedure. The 10-fold CV was finally performed 50 times to avoid the fold randomness bias(83). The Dice coefficient was then calculated between the 10-fold-out group atlas (90 or 91-subjects group-level atlas) and the MWMA.

#### Verification of atlas boundaries

To support the validity of the MWMA, we examined the MRI features and neurotransmitter similarity transitions. Each voxel’s homogeneity was assessed by computing the average of the cosine similarities of MRI features/neurotransmitter between this voxel and all voxels within its local area. This voxel’s local area was centered at the voxel with a radius of five voxels (10 mm).

The macro-level analysis measured cosine similarities across six MRI features for each voxel, including ALFF, dALFF, ReHo, DC, FA, and MD. For each WM voxel of each subject, we calculated its feature homogeneity, i.e., average similarity within the voxel’s local area. Then, the feature homogeneity images were averaged across subjects. The number of distinct ROIs in the voxel’s local area was counted based on the MWMA. Subsequently, Spearman’s rank correlation between the group averaged feature homogeneity, and the number of distinct ROIs across voxels was evaluated. To avoid bias due to spatial autocorrelation, the statistical significance of the correlation result was determined using the Moran spectral randomization (MSR) test, for a total of 5,000 times(84).

In molecular level validation analyses, the neurotransmitter cosine similarity matrix was first measured across nine receptors/transporters density mean images, including ɑ_4_β_2_, CB_1_, D_2_, DAT, GABA_A/BZ_, H_3_, M_1_, mGluR_5,_ and VAChT, from a freely avaliable dataset that integrates group-average receptors/transporters density images from different PET studies(59). Receptors/transporters with more than one mean image, including mGluR_5_ and VAChT, were combined by subjects-weighted averaging. Then, the receptor images were resampled to 2 x 2 x2 mm^3^ to match the resolution of the T1w images. Before the calculation of similarities, each neurotransmitter receptor density data was z-scored across WM voxels. Spearman’s rank correlations between receptor homogeneity and the number of distinct ROIs across all WM voxels were then evaluated. The statistical significance of the correlation result was determined by the MSR, for a total of 5,000 times.

In addition, we computed the network-based random atlases to generate null models. A trivial network was generated by setting all edges between each voxel and its 26-nearest neighbors to one, which preserved the empirical structure of the individual similarity network. A random network was then computed by swapping each edge in the trivial network twice while preserving the degree distribution and connectivity using the Brain Connectivity Toolbox(85). Then, spectral clustering was applied to the random networks. This procedure was repeated 200 times to minimize the random error on the edge swap. Finally, the relationship between MRI features/neurotransmitter homogeneity and the number of distinct ROIs across all WM voxels was calculated based on random atlases, to generate the null distributions.

### Comparison with the existing tract-based JHU-DTI atlas

To measure whether the MWMA better captured functional and topological organizations of WM, we compared local and global metrices between the MWMA and the JHU-DTI-WMPM Type-I atlas. To improve the signal-to-noise ratio, we smoothed the functional images by averaging the time series in a 3-dimensional spherical area with a radius of 10 mm(86).

#### Local evaluation by the DCBC

The DCBC was used to estimate the functional validity of parcellation boundaries from the local perspective(60, 61). The DCBC was computed as the difference between the averaged FC between voxels within the same ROI and the averaged FC between voxels in different ROIs at a given spatial distance. A positive DCBC meant that the boundary of the atlas accurately captured the functional boundaries, and a high DCBC meant high within-ROI homogeneity and between-ROI heterogeneity. In this study, we computed the DCBC between the voxel pairs at a Euclidean distance from 2 to 20 mm, with a step of 2 mm. Then, we compared the DCBC values of MWMA and JHU-DTI atlas at all distances by paired t-test, and corrected by false-discovery rate (FDR) analyses.

#### Global evaluation by the dispersion of functional gradients

Gradients provide a global view of functional organization. The group FC matrix was obtained by averaging individual FC matrices based on the MWMA and the tract-based JHU-DTI atlas. Then, the group FC matrix was row-wise thresholded, i.e., the top 10% of edges in each row were maintained based on our previous study(48, 73). The Gaussian kernel was used to yield the affinity matrix by calculating the similarity of each paired region, with the inverse kernel width set at 1/n, and n was equal to the number of ROIs contained in each atlas, that is, 200 for MWMA and 101 for JHU-DTI atlas (101 of the 102 ROIs of the JHU-DTI atlas were in our group-level WM mask). Next, diffusion map embedding was performed to derive gradients, using the BrainSpace toolbox(84). The embedding algorithm was controlled with two parameters, ɑ and t. We set ɑ at 0.5 and t at 0, which were settings that maintained the global relationship between data points in the embedded space.

In our investigation, the first two gradients were computed for analysis. To better interpret the spatial pattern of the gradient, we projected the gradient points into functional networks, and sorted the first gradient scores within a functional network. Then, we estimated the dispersion of functional gradients at network and global levels(63). The within-network dispersion was quantified as the Euclidean distance of each network node to the corresponding network centroid (centroid of nodes within a network). The global dispersion was quantified as the Euclidean distance of each node to the global centroid (centroid of all nodes).

### Mapping the FC pattern of CC subregions

To ensure the applicability of the MWMA in WM functional analysis, we mapped the FC pattern of identified subregions of the CC. Both the ROI-level and voxel-level FC were calculated. The time series of ROIs were extracted by averaging the signals of all voxels within each ROI, and then z-scoring them. For the ROI-level FC, Pearson’s correlation coefficients between the time series of seed ROI and other ROIs were estimated, resulting in an ROI-level FC (200 x 1) for each subject. For the voxel-level FC, Pearson’s correlation coefficients between the time series of seed ROI and all voxels were estimated, resulting in a voxel-level FC (50,356 x 1) for each subject. Fisher’s r-to-z transformation was performed for individual FC matrices to obtain individual z matrices, then an inverse Fisher’s r-to-z transformation was performed after obtaining the average of individual z matrices.

The reproducibility of the FC pattern was defined as the spatial similarity between the ROI-level and voxel-level FC. To ensure the same dimensionality, the voxel-level FC was downsampled by averaging the FC value of voxels within each ROI to obtain a downsampled voxel-level FC. Before calculating the spatial similarity, the autocorrelation of seed ROI was discarded. Then, the Pearson’s correlation coefficient was calculated between the ROI-level FC (199 x 1) and downsampled voxel-level FC (199 x 1) to measure the spatial similarity. The statistical significance of the correlation result was determined by the MSR test, 5,000 times, and then corrected by FDR analyses.

## Acknowledgments

This work was supported by the National Key Project of Research and Development of Ministry of Science and Technology (2022YFC2009906 and 2022YFC2009900), the National Natural Science Foundation of China (82202250, 62036003, and U1808204), and the Fundamental Research Funds for the Central Universities (ZYGX2022YGRH008). Research reported in this publication was supported by grants1U54MH091657 and by the 16 NIH Institutes and Centers that support the NIH Blueprint for Neuroscience Research; by the McDonnell Center for Systems Neuroscience at Washington University.

## Declaration of interests

The authors report no biomedical financial interests or potential conflicts of interests.

## Data availability

Structural and functional neuroimaging data of normally participants f were obtained through the Washington University-University of Minnesota (WU-Minn) Human Connectome Project (https://www.humanconnectome.org/software/connectomedb). PET-derived tracer images were obtained in neuromaps (https://github.com/netneurolab/neuromaps). The derived MWMA can be obtained from https://github.com/weiliao81/MWMA.

## Code availability

The neuroimaging preprocessing software is freely available (FSL v6.0, https://fsl.fmrib.ox.ac.uk/fsl/fslwiki). The column-pivoted QR factorization (CPQR) algorithm was from https://github.com/asdamle/QR-spectral-clustering. Gradient analysis was used the BrainSpace toolbox (https://brainspace.readthedocs.io). The code used in this study was available at https://github.com/weiliao81/MWMA.

## Notes

### Competing Interest Statement

The authors have declared no competing interest.

