## Supplemental Methods for "Multimodal, multifaceted Imaging-based Human Brain White Matter Atlas"

Number of Figure(s): 6

Supporting Material (s): 1

### Supplementary Methods

***Spontaneous fluctuation*.** The amplitude of low frequency fluctuations (ALFF) reflects the power of spontaneous fluctuation. ALFF was computed as the mean square root of the power spectrum's frequency between 0.01-0.1 Hz in the BOLD-fMRI time series(1).

***Dynamic fluctuation*.** To capture the dynamic spontaneous fluctuations, we measured the dynamic ALFF (dALFF) using a sliding-window approach based on our previous studies(2, 3). In this study, a sliding-window length of 180 TRs was selected to balance capturing the dynamic brain activity and obtaining the reliable estimates of brain activity(4). The sliding window was systematically shifted with a step size of 180 TRs, which means that there is no overlap between the windows. Therefore, ten sliding-windows produced by this procedure. For each sliding window, ALFF was calculated for every voxel in WM mask. The temporal variability of dALFF for each voxel was calculated as the variance of dALFF across the ten sliding-windows.

***Regional homogeneity.*** The regional homogeneity (ReHo) was defined as Kendall's concordance coefficient of a given voxel’s BOLD-fMRI time series with its nearest neighboring voxels’ BOLD-fMRI time series, here, 26 voxels were defined as the nearest neighbors(5).

***Degree centrality*.** The degree centrality (DC) can measure the connectivity capacity of an entire network(6, 7). Individualized FC matrix of WM was firstly constructed by computing the Pearson correlation coefficient between any two given voxels’ BOLD-fMRI time series, resulting in a full matrix (50,356 × 50,356). Then, we generated the thresholded positive network by keeping the positive edges of the connectivity network and setting all the negative edges to zero(8). Next, DC was computed as the column sum of the thresholded FC.

***Structural indices*.** DWI enables the assessment of WM integrity in human brain by providing quantitative measures of FA and MD. The FA effectively measures the degree of anisotropy of a voxel, which is indicative of the direction and organizational structure(9, 10). The MD is another measure obtained from DWI data, and describes the rotationally invariant magnitude of water diffusion within brain tissue(9).

### Supplementary Figures

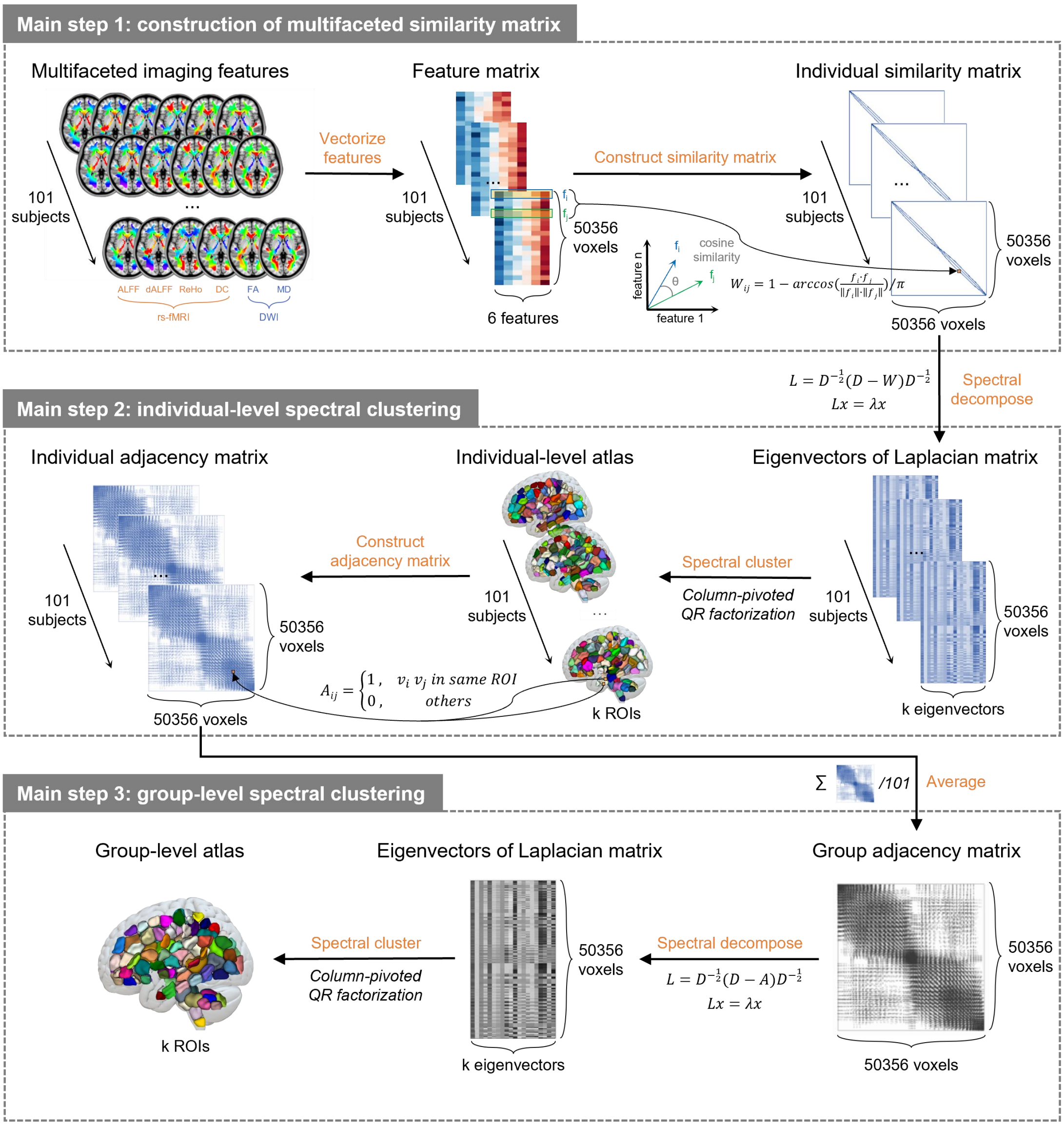


**Supplementary Figure 1. The two-level features similarity network-based clustering parcellation framework.** This parcellation framework was summarized into three main steps, including construction of the multifaceted similarity matrix, individual-level spectral clustering, and group-level spectral clustering. The clustering steps were as follows. Step 1: the feature matrix was vectorized from six multimodal, multifaceted imaging features, including the amplitude of low frequency fluctuations, dynamic amplitude of low frequency fluctuations (ALFF), degree centrality (DC), and regional homogeneity (ReHo), fractional anisotropy (FA) and mean diffusivity (MD) for every subject. Step 2: the individual multifaceted similarity matrix was constructed by calculating cosine similarity of each pair of voxels in the adjacent 26 nearest voxels. Step 3: the individual similarity matrix was converted into the corresponding Laplacian matrix, and spectral decomposition was performed to obtain k eigenvectors corresponding to the k smallest eigenvalues. Step 4: the k eigenvectors were clustered into k regions of interest (ROIs) to complete individual-level spectral clustering. Step 5: the individual adjacency matrix was constructed from individual atlas. Step 6: the individual adjacency matrices were averaged to obtain the group adjacency matrix. Step 7: the group averaged adjacency matrix was converted into a corresponding Laplacian matrix, and spectral decomposition was performed to obtain k eigenvectors corresponding to the k smallest eigenvalues. Step 8: the k eigenvectors were clustered into k ROIs to complete group-level spectral clustering.

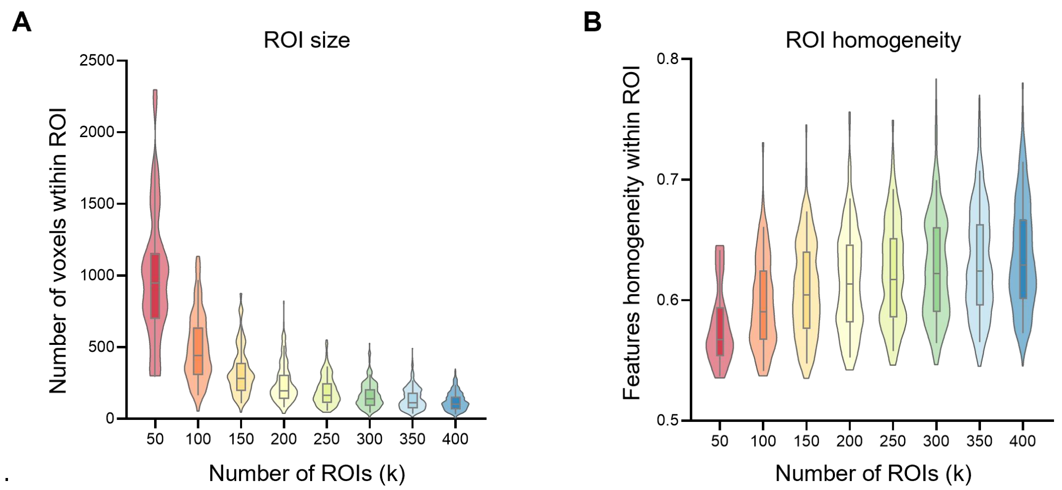


**Supplementary Figure 2. The spatial size and features homogeneity of the region of interest (ROI) in different atlas resolutions.** As the multimodal white matter atlas resolution increased, the mean ROI size and variability of ROI sizes across all ROIs decreased, and the features homogeneity of each ROI increased.

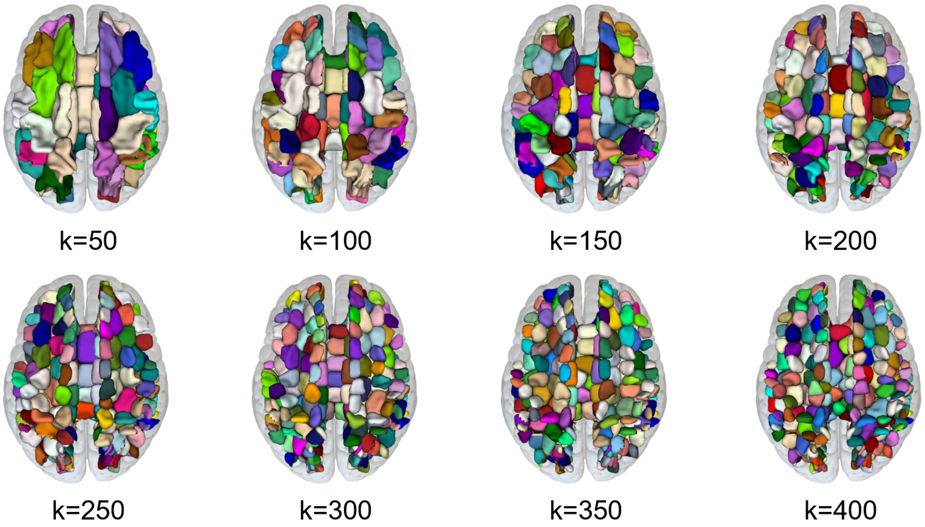


**Supplementary Figure 3. The parcellation pattern of the corpus callosum (CC) in different atlas resolutions.** In the relatively lower resolution from the 50-region of interest (ROI) to 300-ROI atlas, the subregions of the CC were not divided into two hemispheres, whereas in the relatively higher resolution 350-ROI and 400-ROI atlas, the subregions of the CC were divided into two hemispheres.

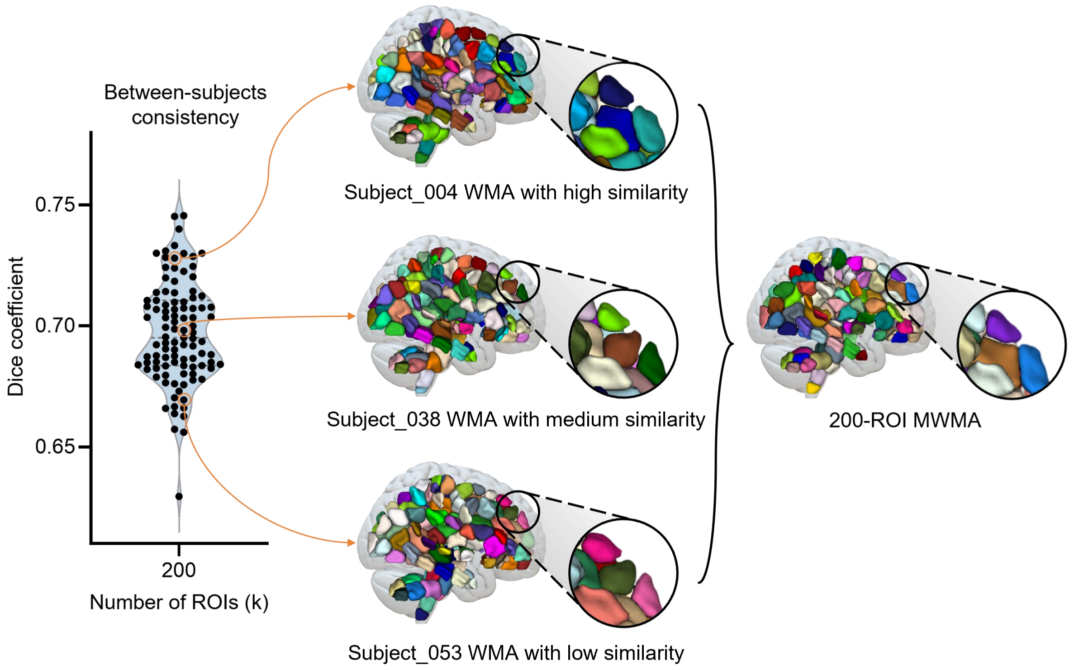


**Supplementary Figure 4. Between-subjects consistency of 200-region of interest (ROI) atlases.** Although there was some variation in the shape or size of different individual atlases, the 200-ROI atlases were substantially similar across subjects, suggesting that this resolution of the multimodal white matter atlas captured the regional arrangement pattern of typical subjects in the healthy young adult population.

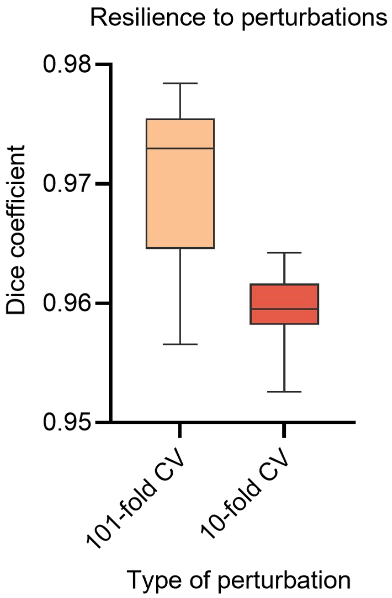


**Supplementary Figure 5. Resilience of 200-region of interest (ROI) multimodal white matter atlas (MWMA) to perturbations.** The 101-fold cross-validation (CV) and 10-fold CV were performed to evaluate the resilience of the 200-ROI MWMA to relatively small and large perturbations. The high-level Dice coefficient of the two types of perturbations indicated the stability of the MWMA.

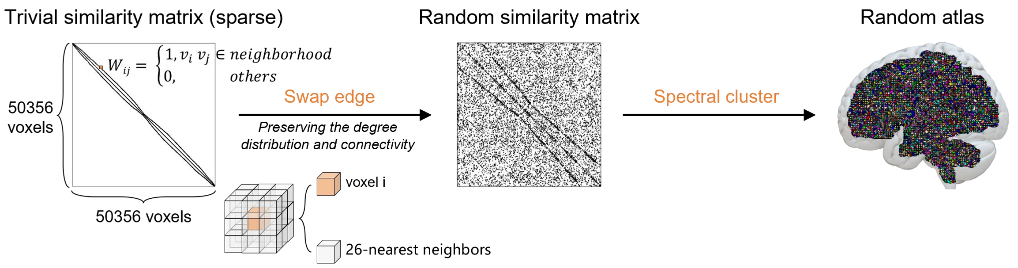


**Supplementary Figure 6. Generation of the network-based 200-region of interest (ROI) random atlases.** To generate the random atlas, we first constructed the trivial similarity matrix, which preserved the network structure of the empirical individual similarity network. Then, each edge of the trivial similarity matrix was swapped twice to generate the random matrix while preserving the degree distribution and connectivity. The random matrix was submitted to the spectral clustering procedure to generate the 200-ROI random atlas. The whole procedure was repeated 200 times to minimize the random error on the edge swap.

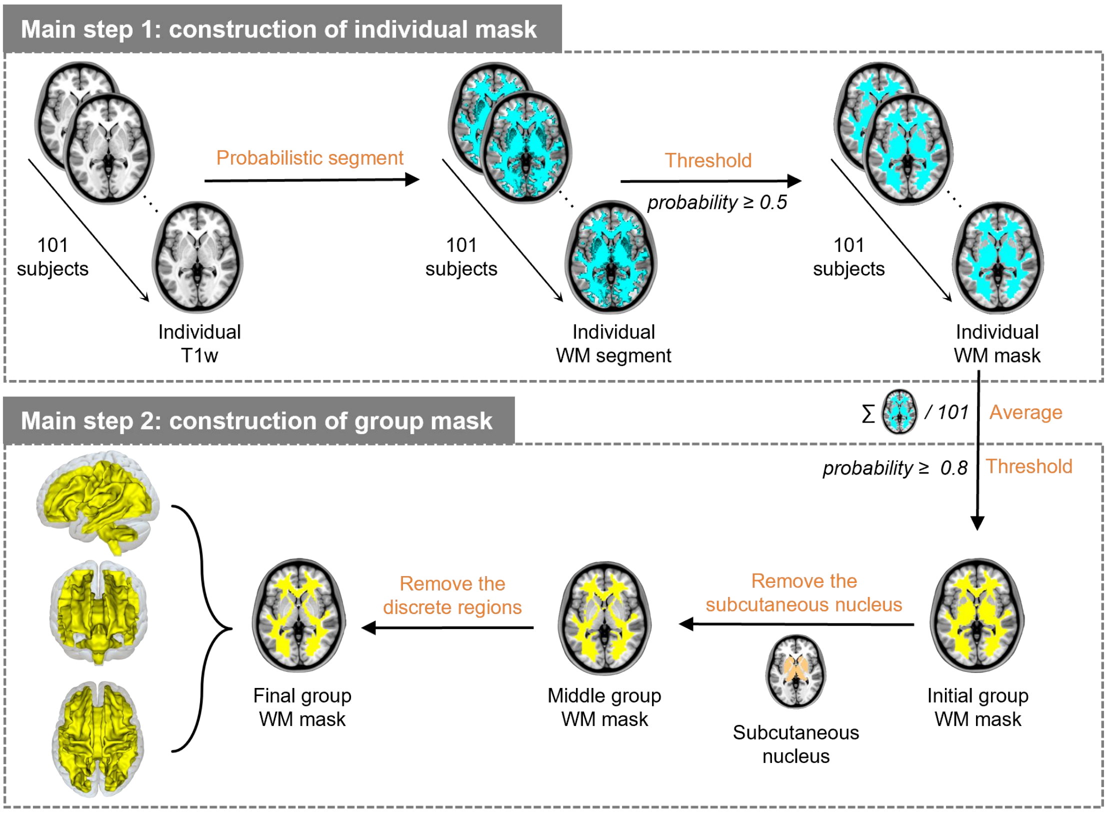


**Supplementary Figure 7. Generation of the group-level white matter (WM) mask.** Step 1: the individual T1w image was probabilistically segmented into grey matter (GM), WM, and cerebral spinal fluid (CSF) for each subject. Step 2: the individual WM mask was created by a threshold of 0.5. Step 3: the initial group-level WM mask was created by averaging individual WM masks and with a threshold of 0.8. Step 4: the middle group-level WM mask was created by removing subcutaneous nuclei from the initial group-level WM mask. Step 5: the final group-level WM mask was created by removing discrete regions on the edge of the middle group-level WM mask, with 50,356 voxels with a resolution of 2 mm.
